# Genotyping Allelic and Copy Number Variation in the Immunoglobulin Heavy Chain Locus

**DOI:** 10.1101/042226

**Authors:** Shishi Luo, Jane A. Yu, Yun S. Song

**Affiliations:** Computer Science Division, University of California, Berkeley, CA, 94720, USA; Department of Statistics, University of California, Berkeley, CA, 94720, USA; Departments of Mathematics and Biology, University of Pennsylvania, Philadelphia, PA, 19104, USA

## Abstract

The study of genomic regions that contain gene copies and structural variation is a major challenge in modern genomics. Unlike variation involving single nucleotide changes, data on the variation of copy number is difficult to collect and few tools exist for analyzing the variation between individuals. The immunoglobulin heavy variable (IGHV) locus, which plays an integral role in the adaptive immune response, is an example of a genomic region that is known to vary in gene copy number. Lack of standard methods to genotype this region prevents it from being included in association studies and is holding back the growing field of antibody repertoire analysis. Here, we establish a convention of representing the locus in terms of a reference panel of operationally distinguishable segments defined by hierarchical clustering. Using this reference set, we develop a pipeline that identifies copy number and allelic variation in the IGHV locus from whole-genome sequencing reads. Tests on simulated reads demonstrate that our approach is feasible and accurate for detecting the presence and absence of gene segments using reads as short as 70 bp. With reads 100 bp and longer, coverage depth can also be used to determine copy number. When applied to a family of European ancestry, our method finds new copy number variants and confirms existing variants. This study paves the way for analyzing population-level patterns of variation in the IGHV locus in larger diverse datasets and for quantitatively handling regions of copy number variation in other structurally varying and complex loci.

## Introduction

The variation between human genomes in gene copy number is understudied and poorly characterized. One such region where this variation is known to exist is the immunoglobulin heavy variable (IGHV) locus. It is a vital component of the adaptive immune system, containing the V genes that code for the heavy chain of antibody molecules. Like other multigene receptor families, the gene segments in this region have accumulated over time through a process of gene duplication and diversification [1–3]. As such, IGHV haplotypes (instances of the IGHV locus) vary not only by single nucleotide polymorphisms, but also in the copy number and ordering of gene segments [4]. However, due to the difficulties of sequencing such complex hypervariable regions, only two full sequences of the locus exist [4,5] and basic population-level characteristics of the locus, such as the relative abundances of different alleles and to what extent individuals differ in copy number, are not known.

The human IGHV locus lies at the telomeric end of chromosome 14 and is approximately 1 Mb in length. Of this 1 Mb region, about 40 functional genes, each approximately 300 bp in length, are used in the biosynthesis of the heavy chain of immunoglobulin (antibody) molecules. There are also approximately 80 non-functional pseudogenes in the region, so-called because they are either truncated or contain premature stop codons. Known allelic variants of individual IGHV genes are currently curated in the International Immunogenetics Information System (IMGT) Repertoire database [6]. The standard nomenclature for IGHV genes is detailed in Materials and Methods.

Given the role of the IGHV locus in the adaptive immune response, IGHV haplotypes are obvious candidates as genetic determinants for susceptibility to infectious disease. Yet due to the lack of a standard format and tools to quantitatively characterize variation in the region, they are not included in association studies. Lack of tools for genotyping the IGHV locus also hampers the burgeoning field of antibody repertoire sequencing [7–11], which is being used in numerous medical applications, including inferring the evolutionary path of broad and potent monoclonal antibodies against human immunodeficiency virus (HIV) [12–14], detecting blood cancers [15, 16], assessing the impact of aging on the antibody response [17], and measuring the adaptive immune response to vaccination [18, 19]. The first step in many of these studies is to align antibody sequencing reads, collected in different individuals, to their germline gene. The current practice is to use germline alleles in a public database of all known alleles (such as the IMGT Repertoire database) for alignment. This is a severe limitation of the process because private IGHV germline configurations can vary substantially between individuals.

Here, we use a high-throughput pipeline to investigate the variation between individuals in the IGHV locus. The pipeline takes reads from whole-genome sequencing (WGS) datasets as input and leverages the IMGT database of known existing alleles. The pipeline performs well on simulated reads from the two known haplotypes (GRCh37 and GRCh38) of this locus. It accurately detects the presence of gene segments from reads as short as 70 bp. With sufficiently long read lengths (250 bp), it also outputs accurate nucleotide sequences of gene segments present in single copy. When we applied the pipeline to genotype the IGHV locus in a sixteen member family, we found evidence of new haplotypes that are mosaics of the existing reference haplotypes and haplotypes that might be transitional between them. We saw examples where offspring inherit structurally different haplotypes from each parent, and where high copy number variation exists within the family. We also identified a putative new allele.

## Results

### Hierarchical clustering to define operational segments

Standard haplotype representations such as single nucleotide polymorphisms and variant call formats are not suitable for genomic regions like the IGHV locus which differ between individuals by insertions and deletions of genes. Given that the functional gene segments of the IGHV locus comprise just 1% of the locus, our approach is to genotype the locus by the copy number and nucleotide sequence of a reference set of functional gene segments.

Defining such a reference set using current IGHV gene nomenclature is problematic, however. IGHV genes that have distinct names can be similar in their nucleotide sequences (Fig. 1A). For example, segments 3-30 and 3-33 in the GRCh37 haplotype should be considered duplicates of the same segment because they differ in only 4 nucleotides (1.4% sequence difference). Indeed, given that some segments have alleles that differ by 5%, determining whether a nucleotide sequence is an allele of segment 3-30 or segment 3-33 is difficult. Across all full-length functional IMGT alleles, there is a significant overlap in the nucleotide differences between alleles with the same segment name and alleles with distinct segment names (Fig. 1B).

**Figure 1.**
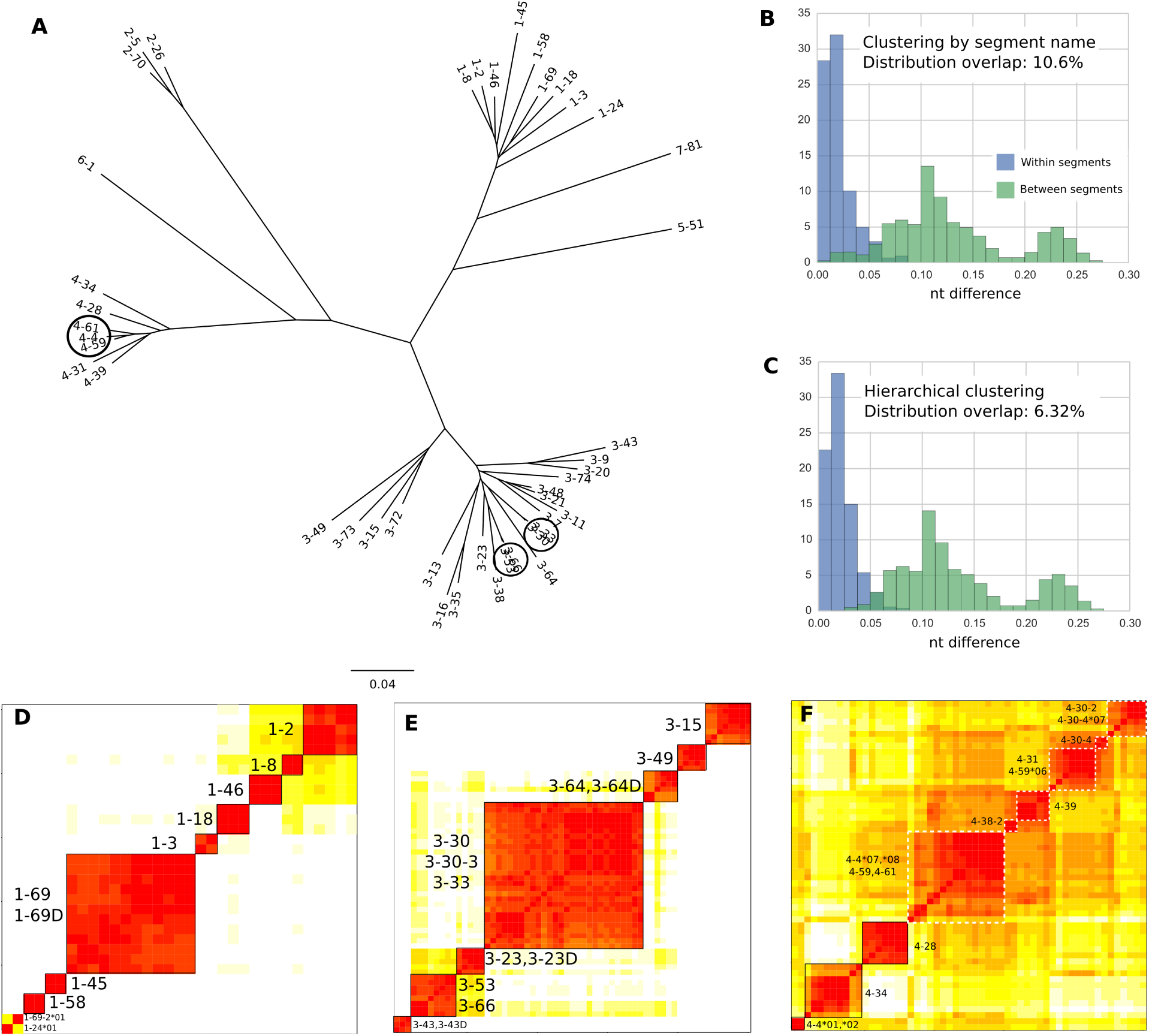
Alleles clustered according to nucleotide similarity. (A) Phylogenetic tree reconstruction of the gene segments in the haplotype sequenced in [5] (which is the GRCh37 haplotype). Circles highlight alleles that are evolutionarily very close. Tree made using neighbor-joining method in ape package in R based on Hamming distance between multiple sequence alignment. (Phylogenetic reconstruction using BEAST [20] led to a qualitatively similar tree). Allele numbers are suppressed for clarity. (B) Distribution of percent nucleotide difference (Hamming distance divided by alignment length) between alleles from same segment (blue) compared against alleles from different segments (green). Alleles from duplicate segments (e.g. 1-69 and 1-69D) have been merged for this analysis. (C) Same as (B) but with alleles partitioned by results of hierarchical clustering rather than segment name. (D-F) Heatmaps of matrices of Hamming distance between alleles. Rows and columns are ordered according to clusters found by hierarchical clustering as described in Materials and Methods. Color spectrum ranges linearly from red to white for nucleotide distances 0-10%. Differences greater than 10% are colored white. (D) Alleles from family 1. (E) Alleles from family 3. Full set of alleles in S3 Fig. (F) Alleles in family 4. Dashed white squares indicate possible clusters.

In order to group alleles according to their sequence similarity, we used hierarchical clustering (Materials and Methods) to identify operationally distinguishable segments. This clustering (Fig. 1C) reduces the overlap compared to clustering by group name alone (Fig. 1B). In family 1, the clusters correspond to segment name, as long as duplicate segments such as 1-69D and 1-69 are merged (Fig. 1D). In family 3, five segments that have distinct names – namely, 3-30, 3-30-3, 3-33, 3-53, and 3-66 – form two clusters {3-30, 3-30-3, 3-33} and {3-53, 3-66} (Fig. 1E). The three segments in family 2 and the two segments in family 5 cluster by segment name (S1 Fig, S2 Fig). Families 6 and 7 each have only one segment and therefore do not require clustering.

Surprisingly, the same clustering algorithm that leads to clean clusters in the other families failed to identify clear-cut clusters in the family 4 set of IGHV genes (Fig. 1F). Not only are the boundaries between clusters fuzzy in this case, but alleles of the same segment cluster separately. For example, 4-4*01 and 4-4*02 cluster separately from 4-4*07 and 4-4*08. The alleles in family 4 also seem to be more similar to each other than alleles in other families. It is not clear why alleles in family 4 in particular should cluster poorly compared to the other families. Gene conversion events in IGHV family 4 and a more recent common ancestor than other IGHV families are both possible explanations that are consistent with the observed distance matrix. A better clustering, based on a combination of evolutionary distance and indel distance, was ultimately used to define the operational segments for family 4 (S4 Fig).

With the caveat that family 4 are more speculative, Table 1 summarizes the operational gene segment clusters defined by hierarchical clustering. Only segments for which the alleles differ from the current IMGT nomenclature are listed. For the remainder of this article, unless stated otherwise, we will use the segment names as they are defined in Table 1.

**Table 1.** Operational gene segments as defined by our hierarchical clustering analysis. Only those that differ from the current IMGT nomenclature are listed.

| Operational name | Alleles, under current IMGT nomenclature |
| --- | --- |
| 1-69 | All 1-69 and 1-69D alleles |
| 2-70 | All 2-70 and 2-70D alleles |
| 3-23 | All 3-23 and 3-23D alleles |
| 3-30 | All 3-30, 3-30-3, and 3-33 alleles |
| 3-43 | All 3-43 and 3-43D alleles |
| 3-53 | All 3-53 and 3-66 alleles |
| 3-64 | All 3-64 and 3-64D alleles |
| 4-4*01 | 4-4*01, 4-4*02 |
| 4-30-2 | All 4-30-2 alleles and 4-30-4*07 |
| 4-31 | 4-30-4*01, 4-30-4*02, 4-31*01-*04, 4-31*10 |
| 4-31*05 | 4-31*05 |
| 4-59 | 4-4*07, 4-4*08, and all 4-59 alleles |
| 4-61 | 4-61*01, 4-61*03-*05, 4-61*08 |
| 4-61*02 | 4-61*02 |

### Pipeline performance on simulated reads

The operational gene segments (Table 1) form the basis of our data pipeline (Fig. 2) that takes whole-genome sequencing reads from an individual and outputs the IGHV genotype (Materials and Methods). We ran the pipeline on simulated reads from the two complete IGHV haplotype sequences (Materials and Methods). Fig. 3 shows the performance of our pipeline at three levels of genotype resolution.

**Figure 2.**
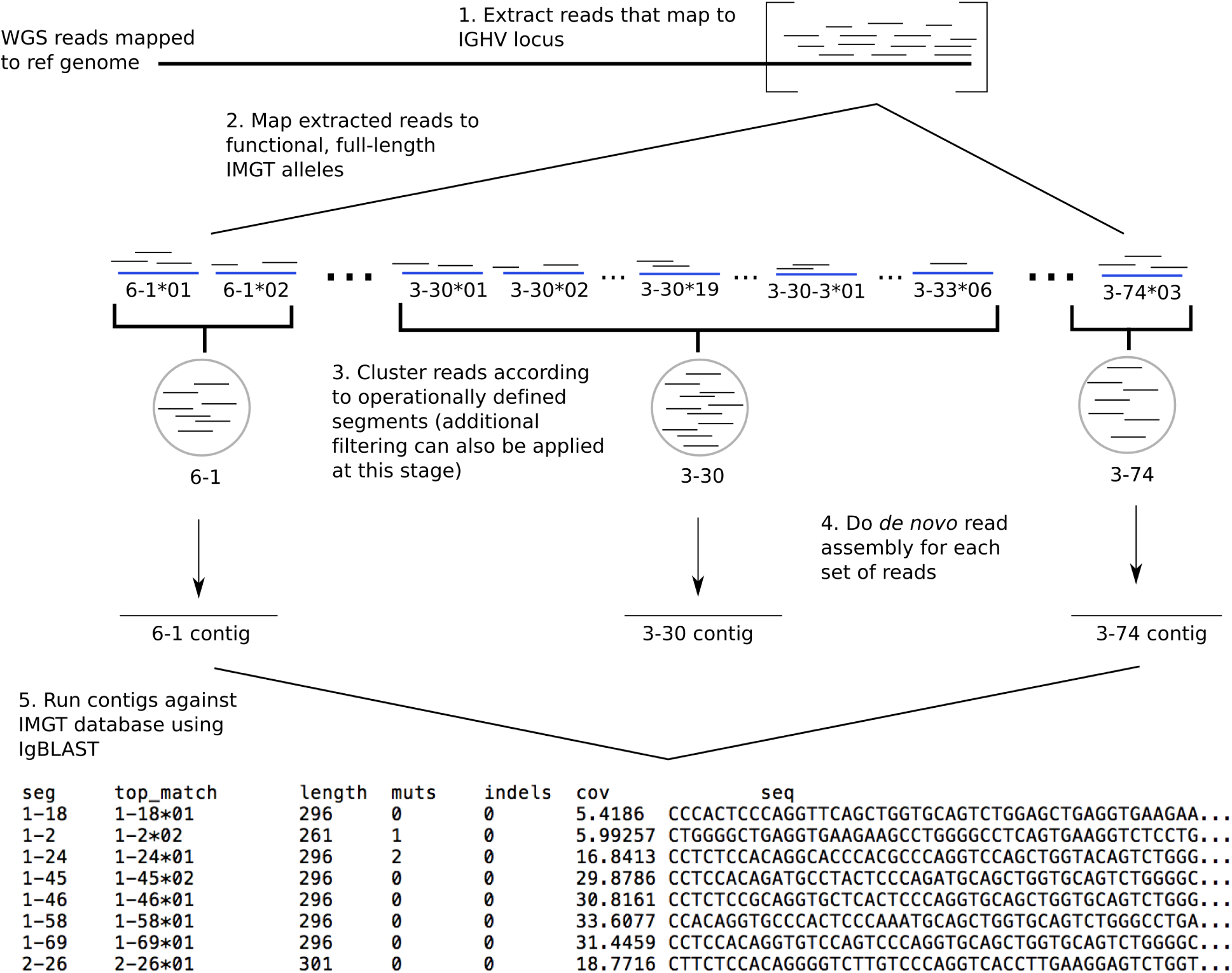
Schematic of genotyping pipeline. 1. WGS reads (short thin black horizontal lines) that map to the IGHV locus of a single individual are extracted from full set of reads. 2. These extracted reads are mapped to known functional IGHV gene segment alleles (thick blue horizontal lines) curated from the IMGT data. 3. Mapped reads are pooled according to the operational segments described in the Results section. At this stage, extra filtering, for example using mate-pair data, can also be applied. 4. Local assembly is performed on reads to produce contigs (long thin black horizontal lines) corresponding to each operational segment. 5. The resulting contigs are identified using stand-alone IgBLAST [21]. The final output contains, for each individual and each assembled contig: the closest-matching existing allele, the length of match, the number of nucleotide mutations or indels that separate the contig from the closest-matching allele, the read coverage of the contig as reported by SPAdes, and the nucleotide sequence of the contig.

**Figure 3.**
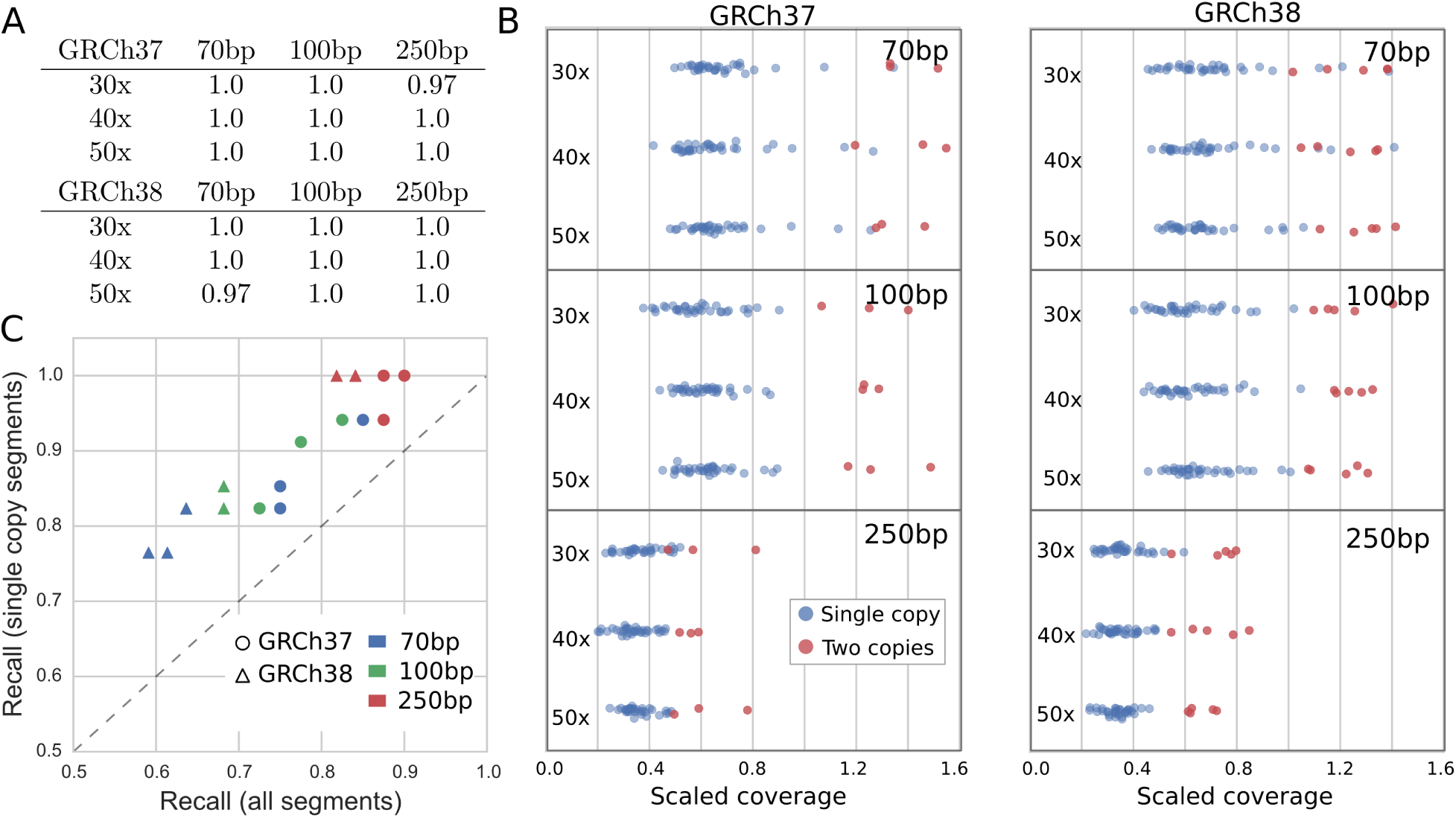
Performance of pipeline on simulated reads from GRCh37 and GRCh38 for varying coverage depths and read lengths. (A) Recall fractions of the pipeline for the two human reference genomes (precision fractions are all 1 and not shown). Recall is calculated as the fraction of operational segments in the reference genome that are correctly called by the pipeline. (B) Read coverage depth of each assembled segment (our proxy for copy number) colored by actual copy number in the reference genome. Raw read coverage depth has been scaled by the simulated coverage depth (e.g. 30, 40, or 50) to allow plots to share same x axis. A slight jitter has been applied to the vertical coordinates of the points to better show their distribution. (C) The recall of alleles for all segments versus the recall for single-copy segments only. Each point is one reference genome, coverage depth, read length combination. Note that two red triangles overlap at the point (0.84, 1.0). Different coverage depths are not indicated as they do not exhibit any pattern.

At the coarsest scale, we ask whether the pipeline correctly identifies the presence or absence of each operational segment. We found the pipeline to be highly accurate, with precision of 100% for all coverage depth (30×, 40×, 50×) and read length (70 bp, 100 bp, 250 bp) combinations and recall of 100% for all but two of the coverage depth/read length combinations (Fig. 3A).

At the next level of resolution, we ask whether the pipeline can correctly determine the copy number of each operational segment. We use the read coverage depth of the assembled contig as a proxy for copy number. Fig. 3B shows that contig coverage depth is indeed correlated with copy number, though some segments which are present in single copy have high coverage depth. This is because pseudogenes in the IGHV locus, which are not included in our reference set, may share common subsequences with functional genes. Reads from pseudogenes can therefore be erroneously mapped, artificially inflating the contig coverage depth. This is particularly an issue with 70 bp length reads as these reads are more likely to completely fall within a conserved region. This problem can be partly alleviated with paired-end reads, a strategy we use on the empirical dataset in the next section.

At the highest level of resolution, we compare the assembled contig obtained from the pipeline to the known nucleotide sequence for each segment. When a segment is only present in single copy in the locus, and if the read lengths are 250 bp, the recall of the segment nucleotide sequence is 100% in all but one of the simulated datasets (Fig. 3C). With shorter reads, the accuracy of correctly calling alleles is lower. As with copy number determination, this lower accuracy is likely due to erroneously mapped reads from pseudogenes that interfere with the assembly algorithm. For the same reason, when a segment is present in more than one copy and as different alleles, the allele calls are also less accurate. Note that higher coverage depth does not necessarily improve accuracy because the error arises not from sequencing error, which occurs in random locations and can be mitigated with higher coverage depth, but from erroneously mapped reads, which are systematically incorrect regardless of coverage.

### Genotyping the Platinum Genomes dataset

We next applied the pipeline to the publicly available Platinum Genomes dataset [22], a set of whole-genome sequencing reads of length 100 bp at roughly 30× depth from a family of 16 (four grandparents, a mother, a father, and ten children, all of European ancestry). Because these reads are paired, we applied an additional filtering step (Materials and Methods) to discard reads that are potentially from pseudogenes in order to improve our allele calls and decrease the false discovery of duplicated genes.

A summary of copy number and allelic variation in IGHV segment types in this dataset is shown in Fig. 4 (S1 Table lists all raw coverage depth values from the dataset). For all the results that follow, the raw coverage depth of each segment has been scaled by the coverage depth of segment 3-74 in the same individual to eliminate variation due to differences in read coverage between individuals. Specifically we assume that 3-74 has two copies, one on each chromosome, and divide the coverage depth of all other segments by half of the coverage depth of 3-74. A normalized coverage depth of 1 therefore corresponds to a single copy on one, but not both, of the chromosomes. Note that the coverage depth tends to decrease towards the 6-1 end of the locus, an issue we will return to in the Discussion.

**Figure 4.**
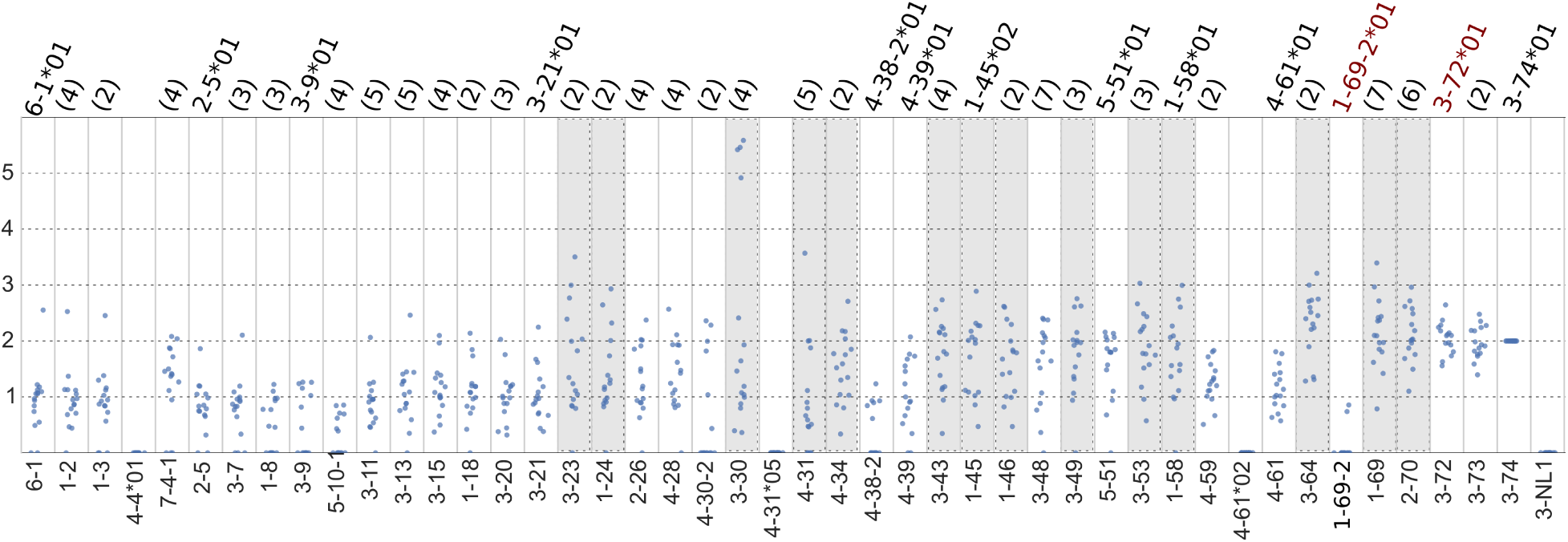
Dotplots of coverage calls for each segment type for Platinum Genomes data. The Y axis is the normalized coverage, i.e. the coverage depth divided by half of the coverage depth of segment type 3-74 (assumed to have two copies). Segment types are ordered, where possible, according to their location in the genome, from 6-1 (centromeric end) to 3-74 (telomeric end). The number in parentheses above each segment type is the number of unique allele sequences found in the family. If only one allele was found, its name is given (in the case where a segment has only one known allele in the IMGT database, the allele name is in red). Shaded columns indicate segment types that likely have more than one copy per chromosome. Note that the outliers for segment types 6-1, 1-2, and 1-3 all correspond to individual NA12891, which had relatively uniform coverage over all segments, unlike other individuals (Fig. 7, S5 Fig). They do not indicate higher copy number. Horizontal jitter has been applied to all points to better illustrate the distribution.

**Figure 7.**
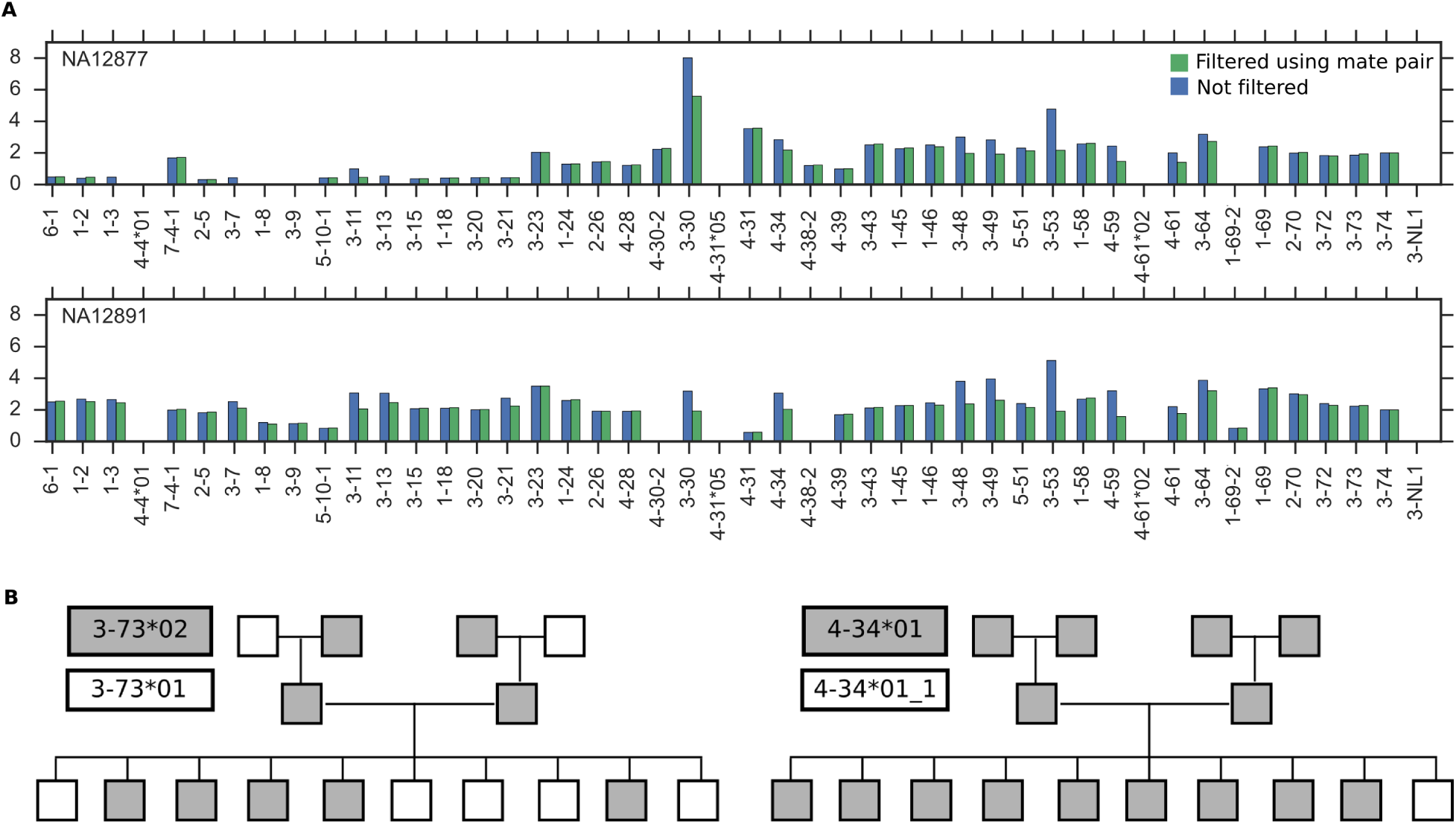
Complications arising from cell type and diploidy in Platinum Genomes dataset. (A) Individuals differ in the uniformity of coverage over segment types when DNA is sequenced from B lymphocytes. Y axis is normalized coverage. (B) Allele calls color-coded by allele and arranged by family tree. For 3-73, at least one of the parents must be heterozygous. For 4-34, the singleton allele in one child indicates that one parent and its parent is heterozygous or that the allele call is incorrect (4-34*01 1 is a variant that is not in the IMGT database and is one nucleotide mutation away from the 4-34*01 allele).

### General patterns of variation

Variation exists in copy number within segments and between segments. Some segments, such as 3-72 and 3-73 are present in all individuals as two copies, one on each chromosome. Others, such as 1-8, 3-9, 5-10-1, 4-38-2, and 1-69-2, are either absent (coverage of zero) or present as a single copy on one chromosome. Coverage depth around the value of three or higher indicates a segment has a duplicate on the same chromosome (segments shaded in grey in Fig. 4). These include 3-23, 3-30, 4-31, 3-43, 3-53, 3-64, 1-69, and 2-70, which are known to have duplicates, but also 1-24, 4-34, 1-45, 1-46, 3-49, and 1-58, for which duplicates have not been previously documented. These latter gene segments are new candidates for copy number variants and topics for further study.

Thirteen out of the forty-two segments (about 30%) found in the family are each represented by the same single allele in all sixteen members of the family. This strongly suggests that these segments are homozygous in the four unrelated grandparents. Genotyping of a larger sample will ascertain whether this set of common alleles is shared for all individuals of European ancestry or is a by-product of our sample being for a small pedigree. In either case, our pipeline and approach begins to address the question of whether a subpopulation can be uniquely identified by a common set of IGHV alleles.

### Multigene copy number variants

We next looked for the presence in family members of two multigene copy number variants that differ between the GRCh37 and GRCh38 reference haplotypes (Fig. 5). Using knowledge of the family pedigree, we are able to reconstruct the diploid genotype for these variants.

**Figure 5.**
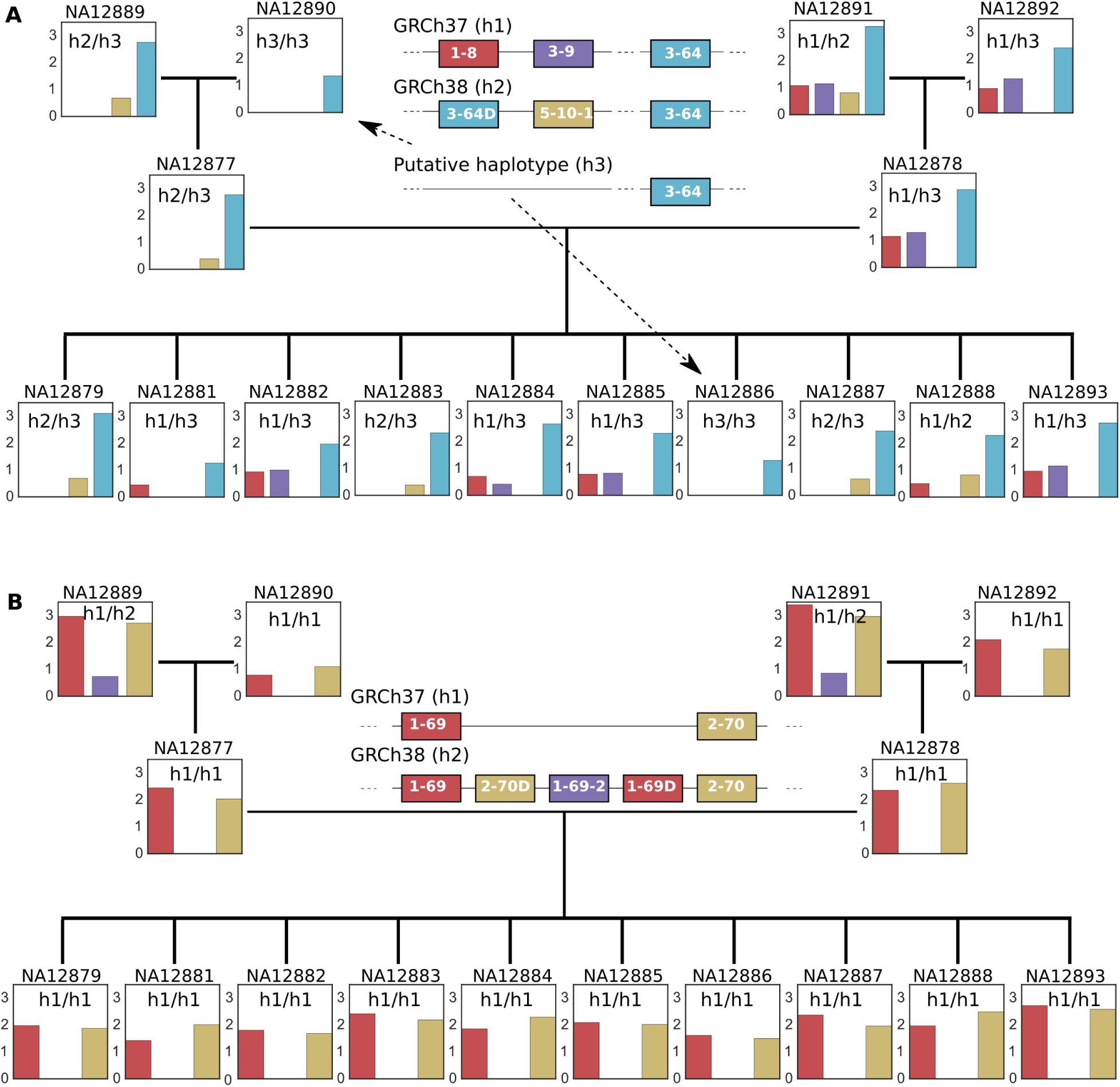
Coverage calls for two multigene copy number variants in GRCh37 and GRCh38. (A) Alternative haplotypes 1-8/3-9 and 3-64D/5-10-1. Additional manual examination of pipeline output shows, consistent with the putative diploid status, that 3-9 is present in NA12881 and NA12888 and that 5-10-1 is not present in NA12886. (B) The insertion haplotype 2-70D/1-69-2/1-69D. In both subfigures, Y axis on each bar plot is normalized coverage. Each bar is colored according to the segment it corresponds to. The putative diploid status for each individual is indicated on the coverage bar plot. Individuals are arranged according to their family tree. NA12877 and NA12878 are the father and mother respectively.

In the case of alternative haplotypes 1-8/3-9 (GRCh37) and 3-64D/5-10-1 (GRCh38), our coverage depth values show that maternal grandparent NA12891 carries both configurations, one on each chromosome. In contrast, the 1-8/3-9 type is entirely absent from the paternal side of the family (Fig. 5A). We manually checked our pipeline output to verify that the copy number calls in the children are consistent with the pedigree. Indeed, both NA12881 and NA12888 which appear to be missing the 3-9 segment, generated reads that mapped to a full-length 3-9*01 allele. This is consistent with NA12881 carrying the GRCh37, but not GRCh38, configuration and NA12888 carrying both the GRCh37 and GRCh38 configurations. (Our automated pipeline did not call the 3-9 segment because the coverage of that segment was too low for the assembler to run). We also verified that NA12890 and NA12886 do not carry the 5-10-1 segment, suggesting a new haplotype, transitional between GRCh37 and GRCh38, which contains a single 3-64 segment without either the 1-8/3-9 or 3-64D/5-10-1 gene combinations. Further, these two individuals appear to be homozygous for this new haplotype, suggesting that the haplotype is common.

For another multigene copy number variant, the 2-70D/1-69-2/1-69D insertion (on GRCh38 but not on GRCh37), grandparents NA12889 and NA12891 carry the insertion on one chromosome and not on the other (Fig. 5B). The insertion did not transmit to the parents or children, with neither the presence of 1-69-2 or elevated coverage for 1-69 and 2-70 present in those individuals.

Interestingly, although all the children are homozygous for the GRCh37 (1-69/2-70) haplotype without the insertion, three of them have the GRCh38 (3-64D/5-10-1) haplotype on at least one chromosome. This implies that there are IGHV haplotypes different from both reference genomes and that are possibly mosaics of the reference genomes. Analysis of these two variants therefore not only confirms the presence of multigene configurations found in the reference genomes in the Platinum Genomes sample, but also demonstrates that different configurations are present in the same ethnic population and that many more configurations may exist.

### High copy number variation in the 3-30 segment

Among all the segments, 3-30 exhibited the most variation in coverage depth (Fig. 4). Using pedigree information as a constraint, we reconstructed the diploid configuration of copy number for segment 3-30 in each member of the family (Fig. 6). Its abundance ranges from zero to four copies on a chromosome. Copy number variation has previously been demonstrated in this region from direct sequencing of the IGHV locus [4]. What is perhaps unexpected from this new analysis is that such high copy number variation can exist in such a small sample; indeed, even between the two chromosomes within the same individual.

**Figure 6.**
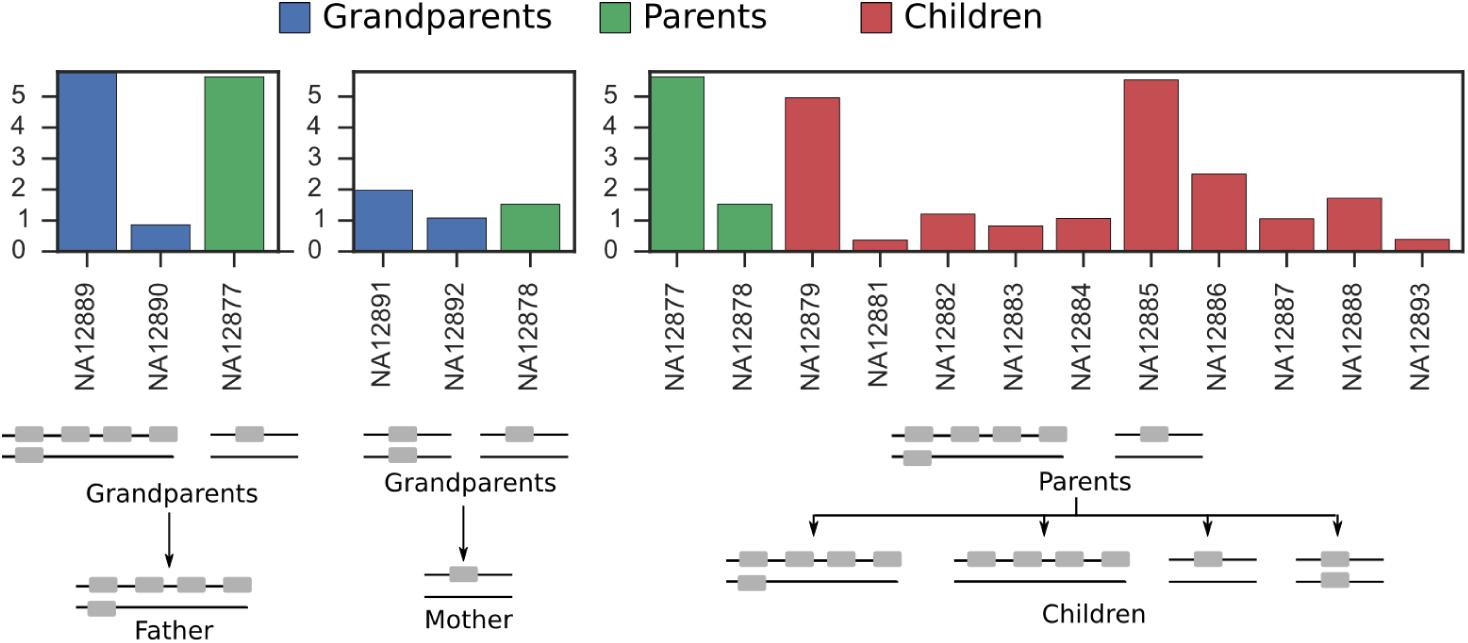
Copy number variation in the 3-30 segment. Below each bar chart of coverage values is a putative reconstruction of the genomic configuration of the individuals in the family. Allele calls are indicated above each bar for reference. Y axis is normalized coverage.

### New 7-4-1 allele

Given the breadth of the IMGT database, we expected to find IMGT alleles amongst allele calls for all the segments in the Platinum Genomes database. This was not the case for segment 7-4-1. The assembled alleles for 7-4-1 all deviated by at least 4 mutations from the nearest IMGT 7-4-1 allele, suggesting the presence of novel alleles for this segment. To eliminate the possibility that the allele calls were confounded by reads from pseudogenes, we applied an extra filtering step to reads that mapped to the 7-4-1 segment (Materials and Methods). A putative new 7-4-1 allele, five nucleotide mutations away from 7-4-1*04, was present in five individuals (S6 Fig shows an alignment of the new allele with 7-4-1*04). The fact that this new allele differs from existing alleles by a large evolutionary distance suggests there are transitional alleles in the 7-4-1 family that are as yet uncharacterized.

## Discussion

With the approach introduced here, we can begin to quantify inter-individual variation in the IGHV locus. Given the small sample size of the Platinum Genomes data, we have focused on quantifying variation in genes known to vary in copy number. As larger whole-genome sequencing datasets become available, it will be possible to compare IGHV copy number haplotypes at the population scale. Machine learning techniques could then be applied to these haplotypes to find correlations between multiple gene segments and discover new copy number variants. Basic open questions such as whether there is a minimal number of IGHV gene segments required for a healthy immune system and whether there is a common core set of IGHV genes that are shared by all individuals can begin to be addressed.

Our study makes clear that read depth information can be used to accurately determine the presence and absence of gene segments. However, complications remain for ascertaining copy number and allelic content to high accuracy. The first complication arises from the cell type on which whole-genome sequencing is commonly performed. The Platinum Genomes data were generated from immortalized B lymphocytes. The IGHV locus in these cell types have undergone VDJ recombination. This rearrangement, which truncates the IGHV locus, confounds the correlation between read coverage depth and copy number of a gene segment. We can see this from the pipeline output, where coverage depth tends to decrease towards the centromeric (6-1 segment) end of the locus. The extent of this decrease can be quite marked, for example in the case of NA12877, or not noticeable at all, for example in NA12891 (Fig. 7A; the distribution of read coverage depth of all the individuals is summarized in S5 Fig). If one knew the number of B cell lineages used to prepare the library and the fraction of haplotypes that underwent rearrangement, it is possible to adjust the raw coverage values to reflect actual coverage values (S1 Appendix). However, in the case of the Platinum Genomes data, this information is unavailable. As whole-genome sequencing becomes more widespread, we anticipate that datasets from other cell types will become available and this issue will be resolved.

The second complication is that the majority of whole-genome sequence reads are generated from diploid cells. Because the majority of segments are on both chromosomes and are of different alleles, the single allele call generated by our pipeline may be composed of sequence from all the alleles present or represent just one of the alleles. Fig. 7B shows that allele calls can hide the heterozygous state of an individual. S7 Fig gives further examples of segments which are present as two alleles in the family and for which the allele calls are misleading. This problem could be addressed with an assembler customized to identify allelic variants of short genomic regions (popular assemblers are currently designed for whole-genome assembly). There has been some success in identifying unique alleles using an alternative data type: antibody repertoire sequencing data [23, 24]. However, such studies cannot directly quantify the copy number of a gene because read abundance in these studies are not correlated with germline gene abundance. Furthermore, the V gene segment is truncated during the genomic rearrangement for producing the antibody coding sequence, so that full-length alleles may not always be obtained from antibody repertoire sequencing data.

We note that there are many existing methods for estimating copy number based on coverage depth using whole-genome sequencing [25–29]. These methods, however, do not utilize the IMGT database of IGHV alleles or specifically target the IGHV locus, a region with a higher amount of repetitions and duplications than most of the genome. They therefore may be prone to biases introduced by targeting the entire genome, which has loci of varying characteristics, rather than targeting a particular region. Additionally, some existing methods [30] intended for whole exome sequencing may be further biased when introduced to data from whole-genome sequencing.

True determination of IGHV haplotypes must ultimately come from sequencing the 1 Mb region in its entirety and in multiple individuals. However, the technology to accurately sequence structurally varying regions remains expensive and low-throughput. We can instead take advantage of the increasing availability of whole-genome sequencing datasets and the extensive IMGT database to genotype this locus in a high-throughput manner. Indeed, not only have we found new haplotypes that are mosaics of reference genome configurations or that are transitional between them with this strategy. The existence of these haplotypes also indicates that our approach of representing the locus in terms of the copy number of a reference set of segments is better suited to cataloging variation in this locus than full sequences of the IGHV locus with annotated breakpoints.

The fundamental strategy applied here is not specific to the IGHV locus. Reads from whole-genome sequencing datasets can similarly be used to characterize other gene families and in other species, where the genes are of comparable length and similar level of diversity. Some examples include T cell receptor genes and olfactory receptor genes. The analysis of whole-genome sequencing data thus need not be restricted to single nucleotide variants, but can also be used to study regions exhibiting copy number variation.

## Materials and Methods

### The standard naming convention of IGHV genes

IGHV genes are named according to their “family” and genomic location. The families, numbered 1 to 7, comprise genetically similar genes. The segment 6-1, for example, is in IGHV family 6 and is the first gene in the locus, counting from the centromeric end. Gene names with a suffix “D” denote a duplicate gene, for example 1-69D, while an appended number, for example 1-69-2, indicates that the gene was discovered subsequent to the original labeling and is located between 1-69 and 2-70. An allelic variant of an IGHV gene is denoted by a *01, *02, etc, as in 1-69*01, 1-69*02.

### Hierarchical clustering

Nucleotide sequences for IGHV gene alleles were downloaded from the IMGT database [31]. Only full-length functional alleles were used for clustering. Multiple sequence alignment was performed on each family of alleles using Fast Statistical Alignment with default parameterization (FSA, [32]). The aligned alleles were then clustered using the hclust function in R (method parameter set to “single”, although using the “complete” method gives the same result for all families with the exception of family 4). For all the IGHV families except family 4, operational segments were determined using distance matrices calculated from Hamming distance based on FSA alignment, with gap differences treated in the same way as mutations. Visual inspection of the alignment of family 4 suggested that indels may be important in partitioning the alleles. Hence, a combination of an evolutionary distance “TN93” (based on [33]) and indel distance (number of sites where there is an indel gap in one sequence and not the other) was used to determine the operational segments for family 4. R scripts are included as a supplementary file (S1 File).

### Genotyping pipeline

Our scripts and example datasets are available at: https://github.com/jyu429/IGHV-genotyping. We assume the WGS data is in bam or sam format [34], with reads already filtered to come from the IGHV locus.

For WGS reads aligned to GRCh37, this is chr14: 105,900,000-107,300,000. For reads aligned to GRCh38, this is chr14:105,700,000-106,900,000 (coordinates extend beyond the IGHV locus to be conservative). Bowtie2 [35] is used to map these reads to all functional, full-length IMGT alleles (the same set used for hierarchical clustering). The default Bowtie2 local alignment threshold led to too many multiple matches. S8 Fig illustrates how we increased this threshold to be more restrictive. Mapped reads are then pooled according to the operational segments described in the Results section. For example, all reads that map to the alleles of 3-30, 3-30-3, 3-33 are pooled together. SPAdes de novo assembler [36] is run on the pooled reads for each operational segment. The assembled contigs are compared with the IMGT database using stand-alone IgBLAST [21] to determine the closest matching allele, the length of match, and the number of nucleotide mutations or indels that separate the contig from the closest-matching allele. The read coverage depth of the contig as reported by SPAdes is also recorded for further analysis.

### Simulated reads

To test the capabilities and quality of our methods, ART [37] was used to generate simulated Illumina reads from GRCh37 and GRCh38 of lengths 70, 100, and 250 bp, each at coverage depths of 30×, 40×, and 50×. Error profiles of simulated reads and adjustments to default ART parameters are illustrated in S9 Fig and S10 Fig.

### Filtering using mate-pair information

For the Platinum Genomes data, which comprises paired-end reads, we apply an additional filtering step to remove reads from pseudogenes that share a common subsequence with a functional gene. One way to disambiguate a read of a functional gene from one of a pseudogene is to compare the genomic position that its mate maps to. If the mate read maps to a region that is substantially farther from the region the first read maps to (we use a threshold of 1000 bp to be conservative) then there is a chance it comes from a pseudogene and the original read is discarded. Note that as a tradeoff, this filtering step will in some cases also incorrectly discard reads from duplicates that are located in a different region of the genome. For segments where the starting position relative to the genome is undetermined, no filtering occurs. In the case of the Platinum Genomes data, which is aligned to GRCH37, this means that filtering is not applied to reads from segments 7-4-1, 5-10-1, 4-38-2, 4-30-2, and 1-69-2.

### Extra filtering step for novel 7-4-1 allele detection

Alleles of 7-4-1 have high nucleotide similarity to subsequences of pseudogenes 7-81, 7-40, and 7-34-1. The mate-pair filtering step above does not apply to 7-4-1 because the Platinum Genomes reads are aligned to GRCh37, which does not contain 7-4-1. To filter out reads from these pseudogenes for 7-4-1, we ran stand-alone IgBLAST on reads mapped to segment 7-4-1. The reads that had the highest match to a pseudogene were removed. The remaining reads were then used as input for SPAdes *de novo* assembler.

## Supporting Information

S1 Fig

Hierarchical clustering applied to Hamming distance between all family 2 alleles.

S2 Fig

Hierarchical clustering applied to Hamming distance between all family 5 alleles.

S3 Fig

Hierarchical clustering applied to Hamming distance between all family 3 alleles.

S4 Fig

Hierarchical clustering applied to all family 4 alleles.

S5 Fig

Dotplots of coverage calls for each individual in the Platinum Genomes dataset.

S6 Fig

Pairwise alignment of the putative 7-4-1 allele, 7-4-1*04 5, with its closest matching IMGT allele, 7-4-1*04.

S7 Fig

Allele calls arranged according to family pedigree.

S8 Fig

Mapped position versus original position of the start of each 70 bp read whose alignment exceeds the score threshold for segment 3-48.

S9 Fig

Error profiles of simulated reads under default ART parameters.

S10 Fig

Error profiles of simulated reads after parameter adjustment.

S1 Appendix

Read coverage adjustment when number of cells is known.

S1 File

R scripts for hierarchical clustering.

S1 Table

Tab-separated (tsv) table of raw coverage depth calls from Platinum Genomes dataset.

## Acknowledgments

The authors would like to thank Erick Matsen and Corey Watson for a number of useful discussions and for their feedback on this manuscript. This research is supported in part by an NSF CAREER Grant DBI-0846015, NIH grant R01-GM094402, and a Packard Fellowship for Science and Engineering.

## References

1. Ota T, Nei M. Divergent evolution and evolution by the birth-and-death process in the immunoglobulin VH gene family. Mol Biol Evol. 1994;11(3):469–482.

2. Niimura Y, Nei M. Evolutionary dynamics of olfactory and other chemosensory receptor genes in vertebrates. J Hum Genet. 2006;51(6):505–517.

3. Das S, Nozawa M, Klein J, Nei M. Evolutionary dynamics of the immunoglobulin heavy chain variable region genes in vertebrates. Immunogenetics. 2008;60(1):47–55.

4. Watson CT, Steinberg KM, Huddleston J, Warren RL, Malig M, Schein J, et al. Complete haplotype sequence of the human immunoglobulin heavy-chain variable, diversity, and joining genes and characterization of allelic and copy-number variation. Am J Hum Genet. 2013;92(4):530–546.

5. Matsuda F, Ishii K, Bourvagnet P, Kuma Ki, Hayashida H, Miyata T, et al. The complete nucleotide sequence of the human immunoglobulin heavy chain variable region locus. J Exp Med. 1998;188:2151–2162.

6. Giudicelli V, Chaume D, Lefranc MP. IMGT/GENE-DB: a comprehensive database for human and mouse immunoglobulin and T cell receptor genes. Nucleic Acids Res. 2005;33(suppl 1):D256–D261.

7. Larimore K, McCormick MW, Robins HS, Greenberg PD. Shaping of human germline IgH repertoires revealed by deep sequencing. J Immunol. 2012;189(6):3221–3230.

8. Robins H. Immunosequencing: applications of immune repertoire deep sequencing. Curr Opin Immunol. 2013;25(5):646–652.

9. Georgiou G, Ippolito GC, Beausang J, Busse CE, Wardemann H, Quake SR. The promise and challenge of high-throughput sequencing of the antibody repertoire. Nat Biotechnol. 2014;32(2):158–168.

10. Calis JJ, Rosenberg BR. Characterizing immune repertoires by high throughput sequencing: strategies and applications. Trends Immunol. 2014;35(12):581–590.

11. Robinson WH. Sequencing the functional antibody repertoire – diagnostic and therapeutic discovery. Nat Rev Rheumatol. 2015;11(3):171–182.

12. Wu X, Zhou T, Zhu J, Zhang B, Georgiev I, Wang C, et al. Focused evolution of HIV-1 neutralizing antibodies revealed by structures and deep sequencing. Science. 2011;333(6049):1593–1602.

13. Liao HX, Lynch R, Zhou T, Gao F, Alam SM, Boyd SD, et al. Co-evolution of a broadly neutralizing HIV-1 antibody and founder virus. Nature. 2013;496(7446):469–476.

14. Doria-Rose NA, Schramm CA, Gorman J, Moore PL, Bhiman JN, DeKosky BJ, et al. Developmental pathway for potent V1V2-directed HIV-neutralizing antibodies. Nature. 2014;509(7498):55–62.

15. Robins H, Desmarais C, Matthis J, Livingston R, Andriesen J, Reijonen H, et al. Ultra-sensitive detection of rare T cell clones. J Immunol Methods. 2012;375(1):14–19.

16. Emerson RO, Sherwood A, Loh ML, Angiolillo A, Kirsch I, Carlson CS, et al. Robust Detection Of Minimal Residual Disease In Unselected Patients With B-Cell Precursor Acute Lymphoblastic Leukemia By High-Throughput Sequencing Of IGH. Blood. 2013;122(21):2550–2550.

17. Jiang N, He J, Weinstein J, Penland L, Saaki S, He X, et al. High Throughput Sequencing of the Human Antibody Repertoire in Response to Influenza Vaccination. J Immunol. 2012;188(Meeting Abstracts 1):58–14.

18. Jiang N, He J, Weinstein JA, Penland L, Sasaki S, He XS, et al. Lineage structure of the human antibody repertoire in response to influenza vaccination. Sci Transl Med. 2013;5(171):171ra19.

19. Jackson KJ, Liu Y, Roskin KM, Glanville J, Hoh RA, Seo K, et al. Human responses to influenza vaccination show seroconversion signatures and convergent antibody rearrangements. Cell Host Microbe. 2014;16(1):105–114.

20. Drummond AJ, Suchard MA, Xie D, Rambaut A. Bayesian phylogenetics with BEAUti and the BEAST 1.7. Mol Biol Evol. 2012;29(8):1969–1973.

21. Ye J, Ma N, Madden TL, Ostell JM. IgBLAST: an immunoglobulin variable domain sequence analysis tool. Nucleic Acids Res. 2013;p. W34–40.

22. Platinum Genomes dataset;. Downloaded 17-member family data sequenced to 50x depth on a HiSeq 2000 system. The reads from one family member, NA12880, failed to index and were not included in analysis. http://www.illumina.com/platinumgenomes/.

23. Boyd SD, Gaëta BA, Jackson KJ, Fire AZ, Marshall EL, Merker JD, et al. Individual variation in the germline Ig gene repertoire inferred from variable region gene rearrangements. J Immunol. 2010;184(12):6986–6992.

24. Gadala-Maria D, Yaari G, Uduman M, Kleinstein SH. Automated analysis of high-throughput B-cell sequencing data reveals a high frequency of novel immunoglobulin V gene segment alleles. Proc Nat Acad Sci. 2015;112(8):E862–E870.

25. Campbell P, Stephens P, Pleasance E, O’Meara S, Li H, Santarius T, et al. Identification of somatically acquired rearrangements in cancer using genome-wide massively parallel paired-end sequencing. Nat Genet. 2008;40:722–729.

26. Alkan C, Kidd JM, Marques-Bonet T, Aksay G, Antonacci F, Hormozdiari F, et al. Personalized copy number and segmental duplication maps using next-generation sequencing. Nat Genet. 2009;41:1061–1067.

27. Chiang D, Getz G, Jaffe D, O’Kelly M, Zhao X, Carter S, et al. High-resolution mapping of copy-number alterations with massively parallel sequencing. Nat Methods. 2009;6:99–103.

28. Yoon S, Xuan Z, Makarov V, Ye K, Sebat J. Sensitive and accurate detection of copy number variants using read depth of coverage. Genome Res. 2009;19:1586–1592.

29. Abyzov A, Urban A, Snyder M, Gerstein M. CNVnator: An approach to discover, genotype, and characterize typical and atypical CNVs from family and population genome sequencing. Genome Res. 2011;21:974–984.

30. Koboldt DC, Zhang Q, Larson DE, Shen D, McLellan MD, Lin L, et al. VarScan 2: Somatic mutation and copy number alteration discovery in cancer by exome sequencing. Genome Res. 2012;22:568–576.

31. IMGT reference directory set;. Accessed: 2014-12-09. Scroll down to ‘IMGT/GENE-DB reference directory sets’. In the table, select Human under IGHV, F+ORF+in-frame P: Nucleotides. http://www.imgt.org/vquest/refseqh.html.

32. Bradley RK, Roberts A, Smoot M, Juvekar S, Do J, Dewey C, et al. Fast statistical alignment. PLoS Comput Biol. 2009;5(5):e1000392.

33. Tamura K, Nei M. Estimation of the number of nucleotide substitutions in the control region of mitochondrial DNA in humans and chimpanzees. Mol Biol Evol. 1993;10(3):512–526.

34. Li H, Handsaker B, Wysoker A, Fennell T, Ruan J, Homer N, et al. The sequence alignment/map format and SAMtools. Bioinformatics. 2009;25(16):2078–2079.

35. Langmead B, Salzberg SL. Fast gapped-read alignment with Bowtie 2. Nat Methods. 2012;9(4):357–359.

36. Bankevich A, Nurk S, Antipov D, Gurevich AA, Dvorkin M, Kulikov AS, et al. SPAdes: a new genome assembly algorithm and its applications to single-cell sequencing. J Comp Biol. 2012;19(5):455–477.

37. Huang W, Li L, Myers JR, Marth GT. ART: a next-generation sequencing read simulator. Bioinformatics. 2012;28(4):593–594.

